# Non-destructive DNA metabarcoding of arthropods using collection medium from passive traps

**DOI:** 10.1101/2023.02.07.527242

**Authors:** Lucas Sire, Paul Schmidt Yáñez, Annie Bézier, Béatrice Courtial, Susan Mbedi, Sarah Sparmann, Laurent Larrieu, Rodolphe Rougerie, Christophe Bouget, Michael T. Monaghan, Elisabeth A. Herniou, Carlos Lopez-Vaamonde

## Abstract

**Background:** Broad-scale monitoring of arthropods is often carried out with passive traps (*e.g*. Malaise traps) that can collect thousands of specimens per sample. The identification of individual specimens requires time and taxonomic expertise, limiting the geographical and temporal scale of research and monitoring studies. DNA metabarcoding of bulk-sample homogenates is faster and has been found to be efficient and reliable, but is destructive and prevents a posteriori validation of species occurrences and/or relative abundances. Non-destructive DNA metabarcoding from the collection medium has been applied in a limited number of studies, but further tests of efficiency are required in a broader range of circumstances to assess the consistency of the method.

**Methods:** We quantified the detection rate of arthropod species when applying non-destructive DNA metabarcoding with a short (127-bp) fragment of mitochondrial COI on two types of passive traps and collection media: 1) water with monopropylene glycol (H_2_O–MPG) used in window-flight traps (WFT, 53 in total); 2) ethanol with monopropylene glycol (EtOH–MPG) used in Malaise traps (MT, 27 in total). We then compared our results with those obtained for the same samples using morphological identification (for WFTs) or destructive metabarcoding of bulk homogenate (for MTs). This comparison was applied as part of a larger study of arthropod species richness in silver fir (*Abies alba*) stands across a range of climate-induced tree dieback levels and forest management strategies.

**Results:** Of the 53 H_2_O-MPG samples from WFTs, 16 produced no metabarcoding results, while the remaining 37 samples yielded 77 arthropod MOTUs in total. None of those MOTUs were shared species with the 389 morphological taxa (343 of which were Coleoptera) obtained from the same traps. Metabarcoding of 26 EtOH–MPG samples from MTs detected more arthropod MOTUs (233) and insect orders (11) than destructive metabarcoding of homogenate (146 MOTUs, 8 orders). Arachnida and Collembola were more diverse in EtOH-MPG samples, but Hymenoptera, Coleoptera and Lepidoptera were less represented than in homogenate. Overall, MOTU richness per trap similar for EtOH–MPG (21.81 MOTUs) than for homogenate (32.4 MOTUs). Arthropod communities from EtOH–MPG and homogenate metabarcoding were relatively distinct, with 162 MOTUs (53%) unique to the collection medium and only 71 MOTUs (23%) present in both treatments. Finally, collection medium did not reveal any significant changes in arthropod richness along a disturbance gradient in silver fir forests. We conclude that DNA metabarcoding of collection medium can be used to complement homogenate metabarcoding in inventories to favour the detection of soft-bodied arthropods like spiders.

## Introduction

Species inventories are a crucial part of ecosystem assessments but are often constrained to a limited number of taxa due to the time-consuming sorting and the need for taxonomic expertise, especially when diverse invertebrate groups are considered (Stork, 2018; Leather, 2018; but see Borkent *et al*. 2018 and Brown *et al*. 2018 who morphologically inventoried dipterans in tropical rainforest). A major breakthrough has been the development of batch-species identification with genetic markers using metabarcoding techniques (Yu *et al*. 2012). Indeed, as this approach identifies species through comparison with DNA barcode reference sequences (Ratnasingham & Hebert, 2007), operators are not required to have taxonomic expertise, providing DNA reference libraries are sufficiently comprehensive and curated by experts (Hebert *et al*. 2003). Despite the incompleteness of DNA reference libraries, metabarcoding has already proven efficient for monitoring arthropod biodiversity (Yu *et al*. 2012), including their response to environmental disturbances (Barsoum *et al*. 2019; Wang *et al*. 2021a; Sire *et al*. 2022).

One major shortfall of the metabarcoding approach is the use of destructive DNA extraction from tissue-homogenate after organisms are dried and ground to fine powder (Yu *et al*. 2012; Sire *et al*. 2022). This prevents the recovery of abundance data and does not allow for a posteriori verification of the specimens, to confirm the presence of a species in a sample. Destructive extraction also prevents further study of the material, such as for integrative taxonomic revisions or even new species descriptions (Marquina *et al*. 2019; Martins *et al*. 2019). Alternative sample preparations have been suggested to facilitate a posteriori morphological control, such as the removal of legs (Braukmann *et al*. 2019) which is time-consuming, or photographing bulk specimens which is a more scalable process but may be insufficient for accurate morphological identification. As for abundance information, optional molecular steps such as DNA spike-in of known mock communities and DNA concentration can also be implemented to infer taxa relative abundance from sequence read-based number correction (Luo *et al*. 2022). Non-destructive DNA extraction buffer (*e.g*. a mixture of lysis buffer with chaotropic salts and proteinase K) has been suggested to keep vouchers intact (Carew *et al*. 2018) and to be suitable for morphological post-examination or DNA re-extraction for confirmatory barcoding (Batovska *et al*. 2021). Although, it was found to be partially destructive after a long incubation time (*e.g*. overnight lysis), especially for soft-bodied taxa like Diptera (Marquina *et al*. 2022; Kirse *et al*. 2022). A recent study also reported the successful attempt of non-destructive DNA extraction from a mix of extraction buffer and propylene glycol acting as preservative solution (Martoni *et al*. 2021). However, these non-destructive alternatives may be limited in terms of scalability by the important volumes and associated costs of extraction buffer required, ranging from 55-65 U.S. $ per Malaise trap sample (Kirse *et al*. 2022).

Shokralla *et al*. (2010) sequenced the DNA of insects from the preservative ethanol (EtOH) solution in which they had been stored (both 40% alcohol mezcal and 95% EtOH preservative solutions). A separate study concluded that DNA metabarcoding of preservative EtOH was a reliable way to identify complex freshwater macroinvertebrate samples (Hajibabaei *et al*. 2012). However, several studies that tried to DNA barcode individual specimens from preservative EtOH reported low amplification success (Robertson *et al*. 2013; Nassuth *et al*. 2014). On the other hand, a study of freshwater arthropod communities using metagenomics of preservative EtOH showed accurate and reliable results, though different from those obtained with shotgun-sequencing of pre-sorted morphospecies of the same samples (Linard *et al*. 2016). In total, 15 other studies have successfully used EtOH-based DNA metabarcoding techniques to characterize complex communities (Zizka *et al*. 2018; Barbato *et al*. 2019; Erdozain *et al*. 2019;; Marquina *et al*. 2019; Gauthier *et al*. 2020; Martins *et al*. 2019, 2020; Milián-Garcίa *et al*. 2020; Young *et al*. 2020; Zenker *et al*. 2020; Couton *et al*. 2021; Persaud *et al*. 2021; Wang *et al*. 2021b; Chimeno *et al*. 2022b; Kirse *et al*. 2022). Most of these studies found dissimilar communities between EtOH-based metabarcoding and their morphological sorting, bulk homogenate or environmental DNA (eDNA) metabarcoding counterparts and highlighted many technical steps to account for those differences. However cross-study comparisons remain difficult as protocols vary in in terms of medium from which DNA is extracted, body structure and size of organisms, primer specificity, bioinformatic pipelines, time prior processing, extraction method (Martins *et al*., 2020). Along with EtOH, there is a growing interest in the applicability of the method on monopropylene glycol (MPG) solutions. Indeed, MPG is widely used for passive traps as it does not attract insects (Bouget *et al*. 2009), is cheaper than EtOH, and evaporates less while preserving specimens. Questions remain regarding the applicability of EtOH, MPG or H_2_O-based metabarcoding in monitoring terrestrial ecosystems, with very few methodological studies focusing on terrestrial arthropods (Marquina *et al*. 2019; Zenker *et al*. 2020; Milián-Garcίa *et al*. 2020; Young *et al*. 2020; Chimeno *et al*. 2022b; Kirse *et al*. 2022).

The present work had three aims: (*i*) comparing the species detected using non-destructive metabarcoding with those detected using either morphological analysis or destructive bulk homogenate metabarcoding, (*ii*) testing the collection medium metabarcoding for two distinct setups commonly used for terrestrial invertebrate biomonitoring, and (*iii*) clarifying the terminology regarding the nature of the medium from which DNA is extracted to facilitate cross-comparability. Finally, we evaluated the impact of forest disturbance levels on arthropod richness to assess the usefulness of non-destructive metabarcoding technique for wide-scale arthropod biodiversity monitoring programs. To do so, we sampled arthropods in silver fir (*Abies alba*)-dominated montane forests along a climate-induced dieback gradient with Malaise trap (MT) and window-flight trap (WFT) setups filled with MPG that was combined with ethanol (EtOH–MPG) and water (H_2_O–MPG), respectively (Figure 1). Metabarcoding of DNA from the collection medium (see Box 1 for terminology) was then compared with the results of different treatments of the same traps: destructive homogenate metabarcoding for MT samples, and morphological identification of Coleoptera to species level for WFT samples (Figure 1).

**Figure 1:**
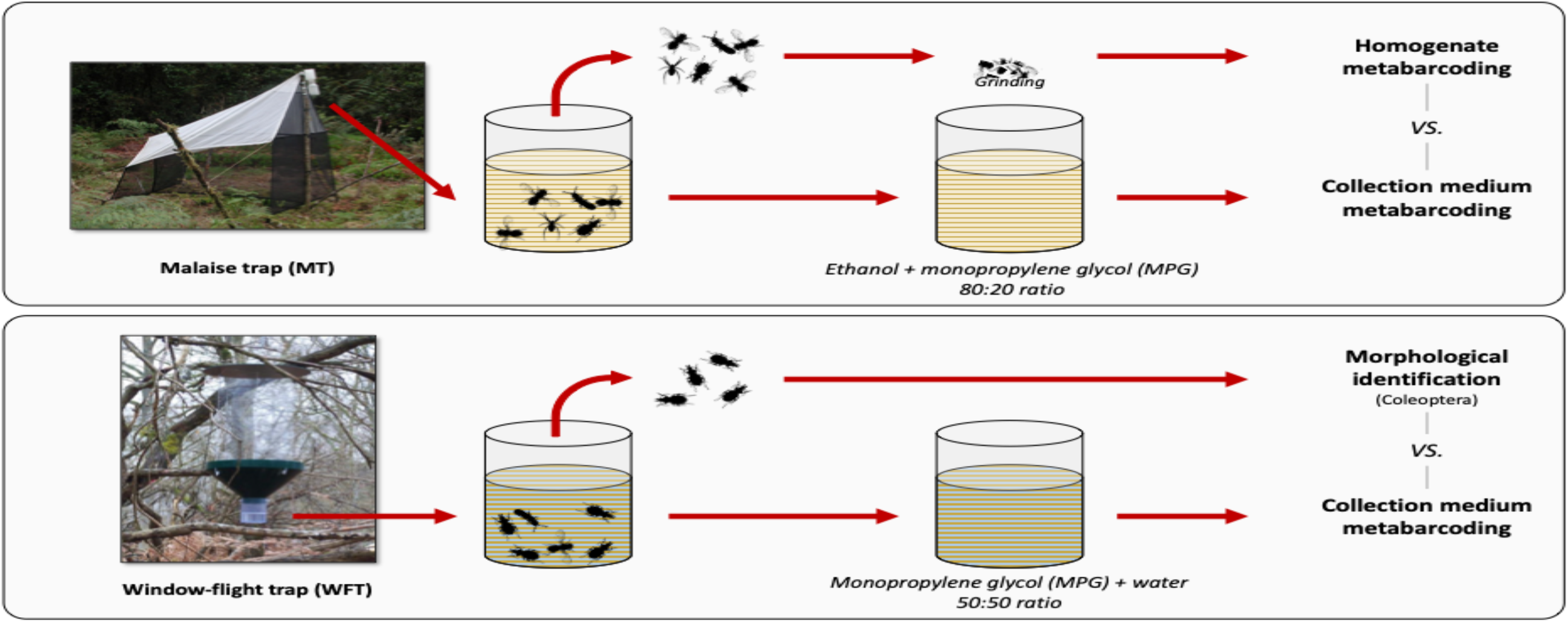
Methodological set-up and sample types processed. Overview of the trapping methods used in this study. For each type of trap, respective collection media (EtOH–MPG for MT and H20–MPG for WFT) are processed through metabarcoding and compared with different treatments (homogenate metabarcoding for MT and morphological identification for WFT) for species detection. All traps were left one month in the field.

## Material & Methods

### Arthropod sampling and environmental assessment

Arthropod communities were sampled between May 15th and June 15th of 2017, in 28 silver fir-dominated forest stands in the French Pyrenees, by following two categorical gradients of climate-induced tree dieback and post-disturbance salvage logging (Sire *et al*. 2022).

In each forest plot, one Malaise trap (MT) was set in the centre with two window-flight traps (WFTs) facing each other at around 10 m-equidistance from it. All traps were left on-site over the entire mid-May to mid-June period. MT collecting jars were filled with ethanol (EtOH) and monopropylene glycol (MPG) in an 80:20 ratio to limit DNA degradation and EtOH evaporation. WFTs were filled with MPG and water (H_2_O) in a 50:50 ratio. After one month in the field, sampling bottles containing the collection medium as well as the arthropods were brought back to the lab and stored in a refrigerator at 4°C for 80 –100 days prior to laboratory processing.

### Arthropod filtration and DNA extraction of homogenate

Arthropods were passively filtered from WFT collection media using single-use coffee filters and were actively filtered from MT collection media using single-use autoclaved cheesecloth and a Laboport^®^ N 86 KT.18 (KNF Neuberger S.A.S., Village-Neuf – France) mini diaphragm vacuum pump connected to a ceramic-glass filtration column that was decontaminated and autoclaved after each use (see Sire *et al*. 2022).

Arthropod bulk filtered from collection media were processed differently for each type of trap (Figure 1). Coleoptera specimens recovered from WFTs were morphologically sorted and identified to species level by expert taxonomists while MT recovered arthropod communities were ground to fine powder using BMT-50-S-M gamma sterile tubes with 10 steel beads (IKA^®^-Werke GmbH & Co KG, Staufen im Breisgau – Germany) and powered at max speed on an IKA^®^ ULTRA-TURRAX^®^ Tube Drive disperser (IKA^®^-Werke GmbH & Co KG). For homogenate metabarcoding from MT samples, DNA extraction was performed from 25 mg (±2 mg) of arthropod powder with Qiagen Dneasy^®^ Blood & Tissue extraction kit (Qiagen, Hilden – Germany) following the manufacturer’s protocol (see Sire *et al*. 2022).

### Filtration and DNA extraction of collection media from MT and WFT samples

Collection medium, as opposed to preservative ethanol, was used throughout the study (see Box 1). Filtration and DNA extraction from collection media were performed for 27 MT (one sample was reported missing) and 53 WFT samples (three samples had technical issues in the field). Sample bottles were agitated by hand for homogenization and filtration was performed by pipetting 100 mL of collection medium with a single-use DNA-free syringe and filtered through a single-use 0.45 μm pore size and 25 mm Ø mixed-cellulose ester (MCE) Whatman^®^ filter (Cytiva Europe GmbH, Freiburg im Breisgau – Germany) held on a 25 mm Ø Swinnex Filter Holder (Merck MgaA, Darmstadt – Germany) that was bleached and autoclaved after each sample filtration. Filters were then placed in DNA-free Petri dishes, cut in half with a sterile scalpel blade and left to dry overnight. After filtering all samples, the filtration step was repeated with molecular grade water to serve as extraction blank control.

DNA extraction from dried filters was done using NucleoSpin™ Forensic Filter kit (Macherey-Nagel GmbH & Co .KG, Düren – Germany). Filter parts were folded and incubated in 600 μL of lysis buffer T1 at 56 °C for two hours with tube horizontally agitated and then centrifugated 1 min 30 sec at 11,000 g. As recommended by Martin *et al*. (2019), we favoured magnetic beads to perform DNA extraction and lysates were processed for DNA extraction using the Macherey-Nagel™ NucleoMag^®^ Tissue kit on an epMotion^®^ 5075vt (Eppendorf, Hamburg – Germany). Volumes on the first binding step were adjusted to the starting volume of lysis buffer accordingly, with 880 μL binding buffer MB2 and 24 μL 0.25X NucleoMag^®^ B-Beads. Extraction was then performed following the manufacturer’s protocol. Final elution was done in 100 μL of elution buffer pre-heated at 56°C with 10 min incubation on beads prior to magnetic separation. Each DNA extraction was quantified using a Qubit^®^ 2.0 fluorometer and the dsDNA High Sensitivity kit (Invitrogen, Waltham (MA) – United States of America).

### PCR amplification of collection media and homogenate

A first but unsuccessful PCR attempt to amplify a 313-bp fragment of the cytochrome c oxidase subunit 1 gene (COI) was performed on collection media using the mlCOIintF (5’-GGWACWGGWTGAACWGTWTAYCCYCC-3’) forward primer and the jgHCO2198 (5’-TAIACYTCIGGRTGICCRAARAAYCA-3’) reverse primer (Leray *et al*. 2013; Geller *et al*. 2013; but see Sire *et al*. (2022) for more details on the PCR conditions).

Successful PCR amplifications to sequence collection media were obtained by targeting a 127-bp fragment of COI using the Uni-MinibarF1 (5’-TCCACTAATCACAARGATATTGGTAC-3’) forward primer and the Uni-MinibarR1 (5’-GAAAATCATAATGAAGGCATGAGC-3’) reverse primer (Meusnier *et al*. 2008). Of note longer 313 bp fragments could not be amplified. Primers were tagged and used in a twin-tagging approach (*i.e*. identical forward and reverse tag for a given sample). The seven bp tags were selected to remain unique after three sequencing mismatches as recommended by Fadrosh *et al*. (2014). No tag was ended in ‘TT’ or ‘GG’ to avoid the succession of three identical nucleotides and potential polymerase slippages. In addition, one to two-bases heterogeneity spacers were added to shift the position of the start of the read and increase nucleotide heterogeneity in the run (Fadrosh *et al*. 2014), and red/green nucleotide balance for Illumina MiSeq technology was checked across all designed tags for increasing nucleotide distinction and sequencing quality (see Supplementary Table I for the full list of tagged-primers).

Before PCR amplification of collection medium DNA samples, qPCR optimization was performed to investigate potential inhibitions and assess the best DNA template dilution. qPCR amplifications were performed with twin-tagged couple #96 of Uni-Minibar primers (see Supplementary Table I; Meusnier *et al*. 2008) on 1/10, 1/20, 1/40, 1/80 and 1/160 serial dilution of DNA template and blank controls in triplicates. qPCR mix was prepared for a 15-μL total volume reaction with 3 μL DNA template, 0.3 μL of each primer (5.5 mM), 7.5 μL of MESA BLUE qPCR 2X MasterMix Plus for SYBR^®^ (Eurogentec) and filled with 3.9 μL of molecular grade water. DNA amplification was performed on a QuantStudio 6 Flex Real-Time PCR System (Life Technologies, Carlsbad (CA) – United States of America) with touch-up cycling conditions as follow: 2 min – 92°C, then 5 cycles of 1 min – 92°C / 1 min – 46°C / 30 sec – 72°C, followed by 35 cycles of 1 min – 92°C / 1 min – 53°C / 30 sec – 72°C before a final elongation step of 5 min at 72°C, as previously described for homogenate DNA, terminated with a high-resolution melting step of 60 sec at 95°C, then 60 sec at 40°C, followed by an acquisition thermal gradient ranging from 65 to 97°C.

Then, the PCR amplifications of collection media samples were run in a 20-μL total reaction volume composed of 5 μL of 1/80 diluted DNA template, 0.2 μL Diamond Taq^®^ DNA polymerase (5.5 U/μL) (Eurogentec, Seraing – Belgium), 2 μL of Buffer (10X) and 3 μL of MgCl2 (25 mM), 0.3 μL of each Uni-Minibar tagged primers (5.5 mM), 0.6 μL dTNPs (20 mM) and filled with 8.6 μL of molecular grade water. PCR cycles were identical as for qPCR optimization. All samples were subject to six replicate PCR reactions, each with a unique primer twin-tag combination from #1 to #31, and samples were distributed in six 96-well plates that also included nine PCR blanks, one filter extraction control for each collection medium and two positive controls.

Finally, we also performed similar PCR amplification of the Uni-Minibar 127-bp amplicon for MT homogenate samples, using 3 μL DNA template at 2 ng/μL, 10.6 μl water and 5+25 PCR cycles. A total of three PCR replicates were performed per homogenate DNA sample distributed in three 96-well plates, each with a specific primer twin-tag combination from #1 to #30 (two blanks and one positive control included). As part of the study by Sire *et al*. (2022), these same homogenate samples had also been processed using Leray/Geller primers (Leray *et al*. 2013; Geller *et al*. 2013) targeting a 313-bp fragment of the DNA barcode and their results are also used here for comparison with this different PCR treatment.

### Library preparation and sequencing of metabarcoding samples

Successful PCR amplification was checked for 10 randomly selected samples for both homogenate and collection media; PCR amplification successes were controlled by migrating 5 μL of PCR product on 2% agarose gel. Homogenate and collection media metabarcoding library preparations were done independently. PCR products of the collection medium samples were purified using CleanNGS (GC biotech, Waddinxveen – Netherland) magnetic beads at a ratio of 0.8 μl per 1 μl PCR product. Purified PCR product was quantified on a FLUOstar OPTIMA microplate reader (BMG Labtech, Champigny-sur-Marne – France) with the Quant-iT™ PicoGreen® dsDNA assay kit (Thermo Fisher Scientific, Waltham (MA) – United States of America) following the manufacturer’s protocol. Equimolar pooling of the samples was carried out for each plate. An additional step with magnetic beads (0.9:1) was added to concentrate the pools to a total DNA quantity of 35 ng of purified amplicon in a final volume of 50 μL. For the library preparation of the pools the NEBNext® Ultra™ II DNA Library Prep Kit for Illumina® (New England Biolabs, Ipswich (MA) – United States of America) was used following the manufacturer’s protocol. Adaptors were diluted 10-fold and a clean-up of adaptor-ligated DNA without size selection was performed. The PCR enrichment step used forward and reverse primers that were not already combined and three amplification cycles. Sequencing was done on an Illumina MiSeq platform using V3 600 cycle kits.

### Bioinformatic and statistical analyses

Bioinformatic demultiplexing was performed following the DAMe pipeline (Zepeda-Mendoza *et al*. 2016, as in Sire *et al*. 2022). Various number of PCR replicates were investigated to retain shared MOTUs with minimum two reads in collection medium metabarcoding (*i.e*. in at least 1/6 PCR replicates, standing as additive demultiplexing; or 2/6; 3/6 and 4/6 for conservative demultiplexing). For homogenate metabarcoding two PCR replicates (2/3) with two reads minimum per MOTU were retained to discard singletons.

MOTU clustering was performed using a 98% similarity threshold and taxonomic assignment was performed with BOLD DNA reference database (Ratnasingham & Hebert, 2007) using BOLDigger tool with BOLDigger option (Buchner & Leese, 2020). Therefrom, taxonomy was retained based on the maximum similarity value of the top 20 hits and correction of top hits was then performed based on the BOLD identification API (Buchner & Leese, 2020). MOTUs with identical species-level taxonomic assignment were then merged manually. Comparisons of MOTU consensus sequences between collection medium and homogenate metabarcoding were performed with BLAST+ (Camacho *et al*. 2009). Only samples with >10k reads were retained and considered in further ecological analyses as samples that could be detecting a representative richness for the given trap types.

All statistical analyses were run with R v4.1.0 (R Core Team, 2017) to test for differences in MOTU recovery between collection medium and homogenate metabarcoding. MT homogenate metabarcoding results of 127-bp amplicons from Uni-Minibar primers were also compared with homogenate metabarcoding of 313-bp amplicons of the same traps (Sire *et al*. 2022). Homoscedasticity of variance and normality of data were checked using ‘descdisc’ and ‘fitdist’ functions from the fitdistrplus v1.1-6 package and assessed with Levene test. If data were normally distributed, an anova test was applied, followed when significant by a pairwise *T*-test with Bonferroni correction. If non-parametric analyses were needed, Kruskal-Wallis tests was applied, along with unpaired Wilcoxon rank-sum tests with Bonferroni correction to assess the direction of the significance when needed. Similar analyses were performed to account for the difference in species richness across dieback level gradient and stand types.

## Results

### Sequencing success, demultiplexing and taxonomic assignment

Sequencing of all collection media samples (EtOH–MPG and H_2_O–MPG) resulted in 12,686,324 reads. MOTUs with at least two reads (*i.e*. to remove singletons) were investigated within different demultiplexing thresholds: from additive (MOTUs present in at least 1/6 PCR replicates) to more stringent demultiplexing (MOTUs present in at least 2/6, 3/6 and 4/6 PCR replicates). Reads were found in 2/11 negative controls for the most restrictive demultiplexing threshold (4/6) and in up to 9/11 negative controls for the additive demultiplexing. Throughout the dataset cleaning process, MOTUs found only in negative controls were removed. This filtering towards raw dataset induced between 71.3% reads drop (from 1405 MOTUs and 10,821,027 reads to 1276 MOTUs and 3,104,116 reads) for the 1/6 additive demultiplexing and 15.8% reads drop (from 210 MOTUs and 7,169,549 to 196 MOTUs and 6,037,276 reads) for the 4/6 demultiplexing threshold (Supplementary Table II). Further filtering implied the removal of non-Arthropoda MOTUs, MOTUs with a similarity to reference sequence below 80%, and the merging of MOTUs with identical species identification. These filtering criteria reduced the number of MOTUs from 1276 to 495 for 1/6 PCR replicates threshold, 471 to 267 for 2/6, 294 to 198 for 3/6 and 196 to 146 for 4/6 (Figure 2; Supplementary Table II).

**Figure 2:**
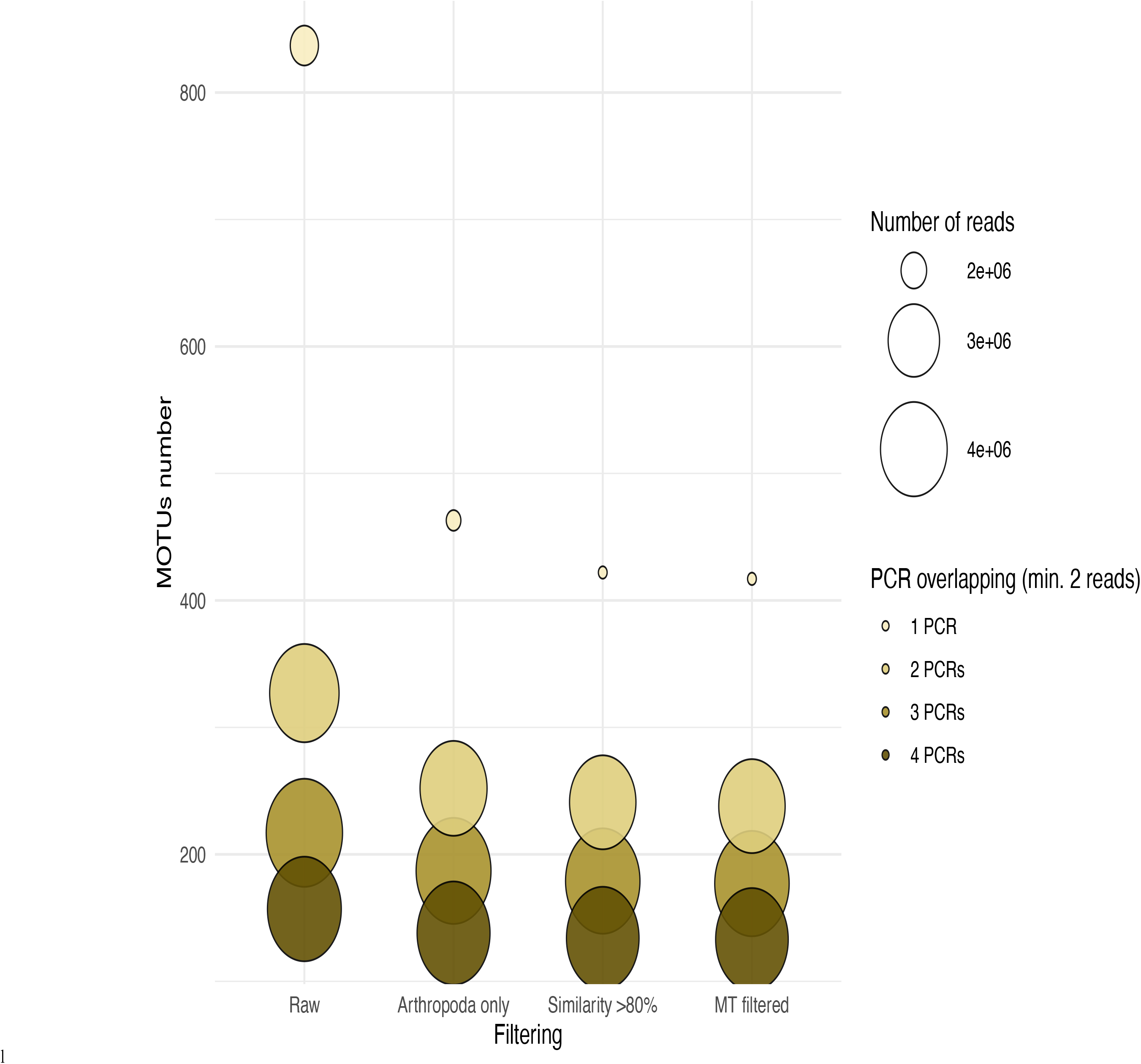
MOTUs and reads numbers after filtering steps of Malaise trap datasets generated with different bioinformatic demultiplexing thresholds. Circles represent the number of MOTUs retained for various filtering and demultiplexing stringency thresholds, with circle wideness corresponding to the associated read numbers. Bioinformatic demultiplexing thresholds are defined by the number of PCR replicates in which a MOTU with a minimum of two reads has to appear to be retained (*i.e*. MOTU present with two reads in at least **1/6** PCR, overlapping **2/6**, **3/6** or **4/6** PCR replicates, coloured from lighter to darker yellow, respectively). Filtering steps are described as follow : **Raw** correspond to the dataset recovered after demultiplexing and removal of MOTUs from blank and positive controls; **Arthropod only** indicates a filtering based on taxonomy to retained MOTUs identified as Arthropods only; **Similarity >80%** corresponds to a filtering based on the percentage of similarity shared with the consensus from BOLD database used for taxonomic identification and keeping MOTUs sharing at least 80% similarity only; **MT filtered** corresponds to the final dataset used for Malaise traps, with a merging of MOTUs and occurrence information based on an identical species identification.

Regarding Window-flight traps (WFTs), 1/6 to 4/6 demultiplexing thresholds of collection medium (H_2_O–MPG) sequencing yielded 191, 77, 53 and 37 MOTUs, respectively, most of them identified as Diptera (100/191, 43/77, 29/53 and 20/37). When focusing on Coleoptera (*i.e*. the main taxonomic group sampled by WFT), only 20/191, 3/77, 2/53 and 2/37 corresponding MOTUs were recovered. In comparison, morphological sorting of the WFT led to 389 morphotaxa, of which 343 species could be identified (Supplementary Table III). A total of 18/20 Coleoptera were identified to species level for the 1/6 demultiplexing threshold. Among these, 12 were also found in the morphological dataset, of which only five were found in the same traps following both metabarcoding and morphology treatments. These observations had low reliability as overall these five species had very few concurrent occurrences among treatments (*i.e*. one sample by metabarcoding out of 13 in morphology, 1/17, 1/17, 1/27 and 3/53, respectively) and multiple detections in metabarcoding samples that were not verified *via* morphological sorting (*e.g*. potential cross-contaminations). Similarly, for the three Coleoptera from 2/6 demultiplexing threshold that were all identified down to species level (*Cis festivus* (Panzer, 1793), *Pyrochroa coccinea* (Linnaeus, 1761) and *Quedius lucidulus* (Erichson, 1839)): *P. coccinea* was not found in the morphological dataset and the other two–also corresponding to the Coleoptera MOTUs found in 3/6 and 4/6 demultiplexing thresholds–were present but not detected concurrently in the morphological and metabarcoding treatments of the same traps (Supplementary Table III, Supplementary Table IV).

For Malaise trap (MT) collection medium, ratios in MOTU reduction from the various filtering steps were similar for all demultiplexing thresholds apart from the additive one (1/6 PCR replicates) which showed a more drastic loss in both reads and MOTUs (Figure 2, Supplementary Table II). We compared 1/6 and 2/6 demultiplexing results to 313-bp bulk metabarcoding results from a previous study on the same MTs (Sire *et al*. 2022). As the two COI fragments were of different length (127 and 313-bp) and did not overlap (Elbrecht *et al*. 2019), we downloaded full-length barcodes of publicly available records matching identification from BOLD for 313-bp derived MOTUs. Comparisons with our 127-bp derived MOTUs from 1/6 and 2/6 demultiplexing thresholds gave only 67 (114 with >97% similarity) and 45 (72 with >97% similarity) identical and shared MOTUs, respectively. Comparing both 127-bp demultiplexing thresholds, 40 MOTUs with 100% similarity to 313-bp dataset were shared. The additional 27 MOTUs from the 1/2 additive demultiplexing are identified as Diptera (16), Lepidoptera (6), Hemiptera (2), Coleoptera (2) and Hymenoptera (1).

While 1/2 demultiplexing threshold allows a slightly better recovery of insects from collection medium metabarcoding of MT samples (*i.e*. 27 additional MOTUs that we could also identify with 313-bp bulk metabarcoding), no improvement was highlighted at that demultiplexing threshold for WFTs. As this led to little increase in MOTUs, and in order to reduce the risks of dealing with false positive MOTUs from 1/6 PCRs threshold, hereafter results focus on the filtered dataset from the 2/6 PCR replicates demultiplexing threshold only. The 27 EtOH–MPG (MT) samples gave a total of 238 arthropod MOTUs and a number ranging from three to 46 (Table I) with 147,358.6 (± 13,687.25 SE) reads per sample. As one trap had <10k reads, it was further removed, giving a final dataset of 233 arthropod MOTUs for 26 successfully metabarcoded samples. Of the 53 H_2_O–MPG (WFT) samples, 37 (70%) yielded arthropod MOTUs for a total number of 77 (Table I; Supplementary Table IV), 12,176.06 (± 5,073.41 SE) reads per sample, with MOTUs number ranging from one to six for all but one sample that harboured 47 MOTUs and a mean of 2.06 MOTUs per sample (Table I). Similar percentages of taxonomic assignment were found for the 233 MOTUs detected in the MT collection medium (EtOH–MPG), 226 (97%) were unambiguously assigned to order, 217 (93%) to family, 145 (62%) to genus and 118 (51%) to species (Figure 3A; Supplementary Table V).

**Figure 3:**
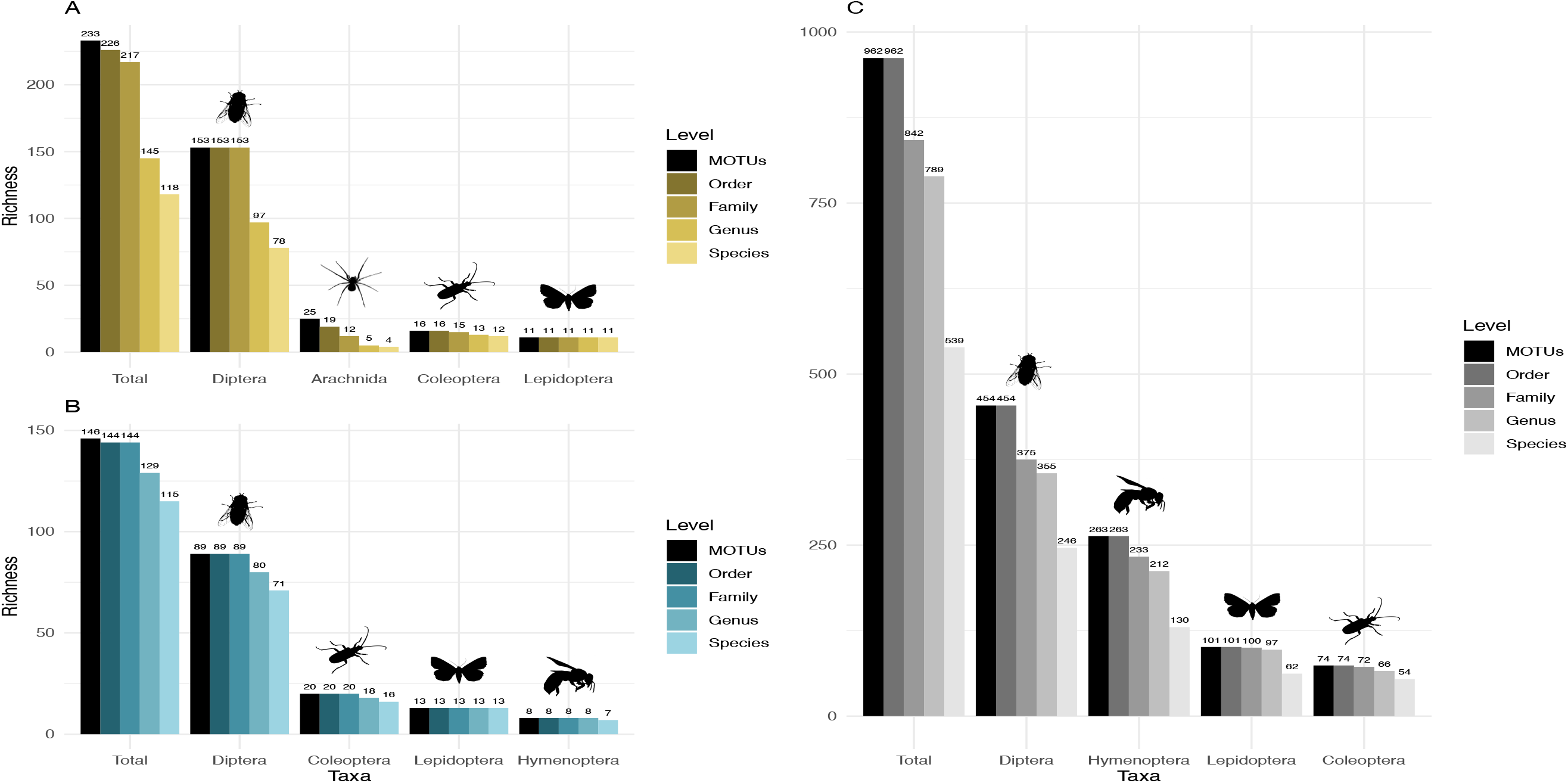
Taxonomic assignment of recovered arthropod MOTUs from collection medium and homogenate metabarcoding of the same Malaise trap samples. Number of MOTUs clustered at 97% similarity from a 127-bp (Uni-Minibar primers) or a 313-bp (Leray/geller primers) COI fragment and taxonomically assigned unambiguously based on BOLD DNA barcode reference libraries. Data are shown for (**A**) collection medium metabarcoding with Uni-Minibar primer set (yellow), (**B**) homogenate metabarcoding with Uni-Minibar primer set (blue) and (**C**) homogenate metabarcoding with Uni-Minibar primer set (gray) of the same Malaise traps. The four most diverse arthropod taxa for each sample type are displayed. Black bars represent the total number of MOTUs for each category and shaded colour gradient bars—from dark to light (yellow, blue or gray) for order to species level, respectively—highlight the number of MOTUs assigned to the associated taxonomic level. Labels provide the number of MOTUs.

Sequencing of MT tissue homogenate targeting the 127-bp amplicon resulted in 3,728,546 reads in total, reduced to 406,776 for 169 MOTUs after applying a demultiplexing threshold of 2/3 PCR replicates with a minimum of two reads per MOTU. Filtering of negative and positive controls generated 75% reads drop (from 406,776 reads to 101,655 for a three MOTUs loss). Two traps yielded no result with homogenate metabarcoding and corresponded to samples with 29 and 46 MOTUs detected in collection medium. Each of the 25 remaining traps harboured one to 50 MOTUs and an average number of reads per sample of 10,982.3 (± 4,139.802 SE). For ecological analyses, 15 traps did not meet the >10k reads threshold and were discarded, leading to a final dataset for homogenate metabarcoding from MT samples comprising 146 arthropod MOTUs for 10 traps (Supplementary Table VI). Taxonomic assignment resulted in 144 (99%) MOTUs assigned to order and to family, 129 (88%) to genus and 115 (79%) to species (Figure 3B). Compared with metabarcoding of the same traps targeting a 313-bp amplicon (Sire *et al*. 2022), our results for a shorter fragment (127-bp) yielded a significantly lower number of MOTUs per trap overall (Wilcoxon rank sum-test: *p* = 1.3^e-05^; Figure 4), as well as across different taxa (Supplementary Figure 1). Further analyses of community diversity only focus on the results of the 127-bp homogenate metabarcoding for comparisons with Malaise trap collection medium metabarcoding using that same shorter fragment.

**Figure 4:**
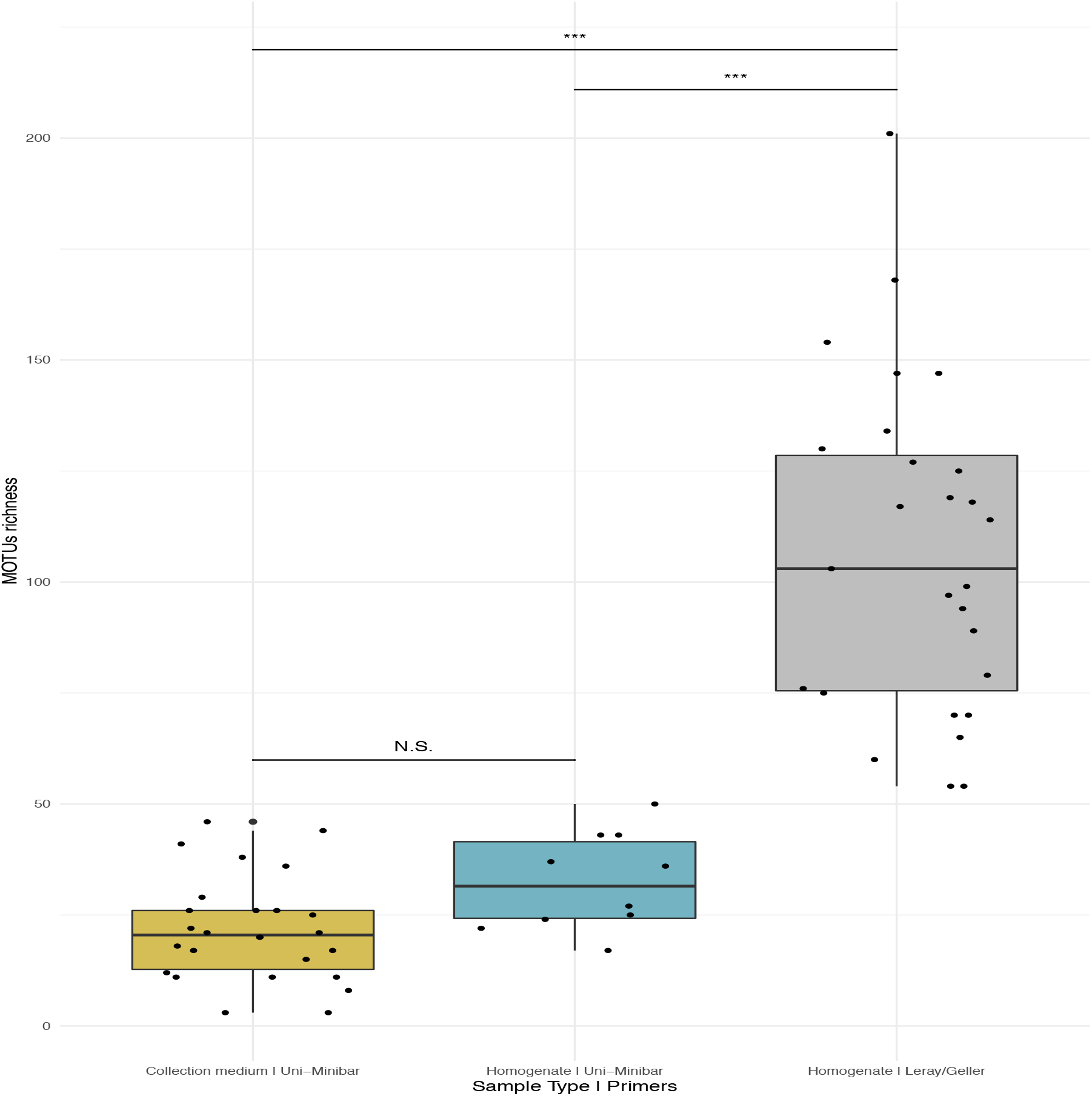
Comparison of MOTU richness recovered from Malaise traps using various metabarcoding treatments (collection medium vs. homogenate) or primer sets (Uni-Minibar vs. Leray/Geller). Boxplot of MOTU count for collection medium (yellow; 1) or homogenate metabarcoding (blue; 2) with Uni-Minibar primer set or from homogenate metabarcoding using mlCOIintF/jgHCO2198 primer set (gray; 3) of the same Malaise trap samples. Black dots represent samples considered after demultiplexing and data curation. Significant differences adjusted with Bonferroni correction are highlighted with ‘*’ and ‘N.S.’ stands as non-significant. Similar MOTU richness could be detected from collection medium and homogenate metabarcoding using Uni-Minibar primers, but significantly lower than the richness detected with a longer amplicon targeted with Leray/Geller primers in a previous study (Wilcoxon rank sum-test: 1–2: *p* = 0.071; 1–3: *p* = 1.3^e-09^; 2–3: *p* = 1.3^e-05^).

### Comparative analyses of community composition between treatments and across forest disturbances

Metabarcoding analyses of WFT collection medium samples yielded only 77 MOTUs, with only three Coleoptera. We focus hereafter on the results from MT samples only. Overall, the MOTUs richness from collection medium metabarcoding (n = 26, mean = 21.80, median = 20.5) was similar than with homogenate metabarcoding (n = 10, mean = 32.4, median = 31.5) (Wilcoxon rank sum-test: 1–2: *p* = 0.071; Figure 4).

Community compositions differed between homogenate and collection medium metabarcoding. Out of the 146 arthropod MOTUs recovered from the MT homogenate, 2% (3 MOTUs) were Collembola, 4% (6 MOTUs) were Arachnida and the remaining 94% (137 MOTUs) were Insecta, while the 233 MOTUs recovered from the MT collection medium were 4% (10 MOTUs) Collembola, 11% (25 MOTUs) Arachnida and 85% (198 MOTUs) Insecta (Figure 5A, B). Insects recovered from EtOH–MPG collection medium belonged to 11 orders: 77% (153 MOTUs) were Diptera, 8% (16 MOTUs) Coleoptera, 6% (11 MOTUs) Lepidoptera, 3% (5 MOTUs) Hymenoptera, and the remaining 7% (13 MOTUs) belonged to Ephemeroptera, Mecoptera, Neuroptera, Psocodea, Raphidioptera, Thysanoptera or Trichoptera (Figure 5C). The insect community from homogenate was composed of eight insect orders and a different distribution MOTUs: 65% (89 MOTUs) Diptera, 15% (20 MOTUs) Coleoptera, 10% (13 MOTUs) Lepidoptera, 6% (8 MOTUs) Hymenoptera and the remaining 5% (7 MOTUs) belonged to Hemiptera, Neuroptera, Psocodea or Raphidioptera (Figure 5D).

**Figure 5:**
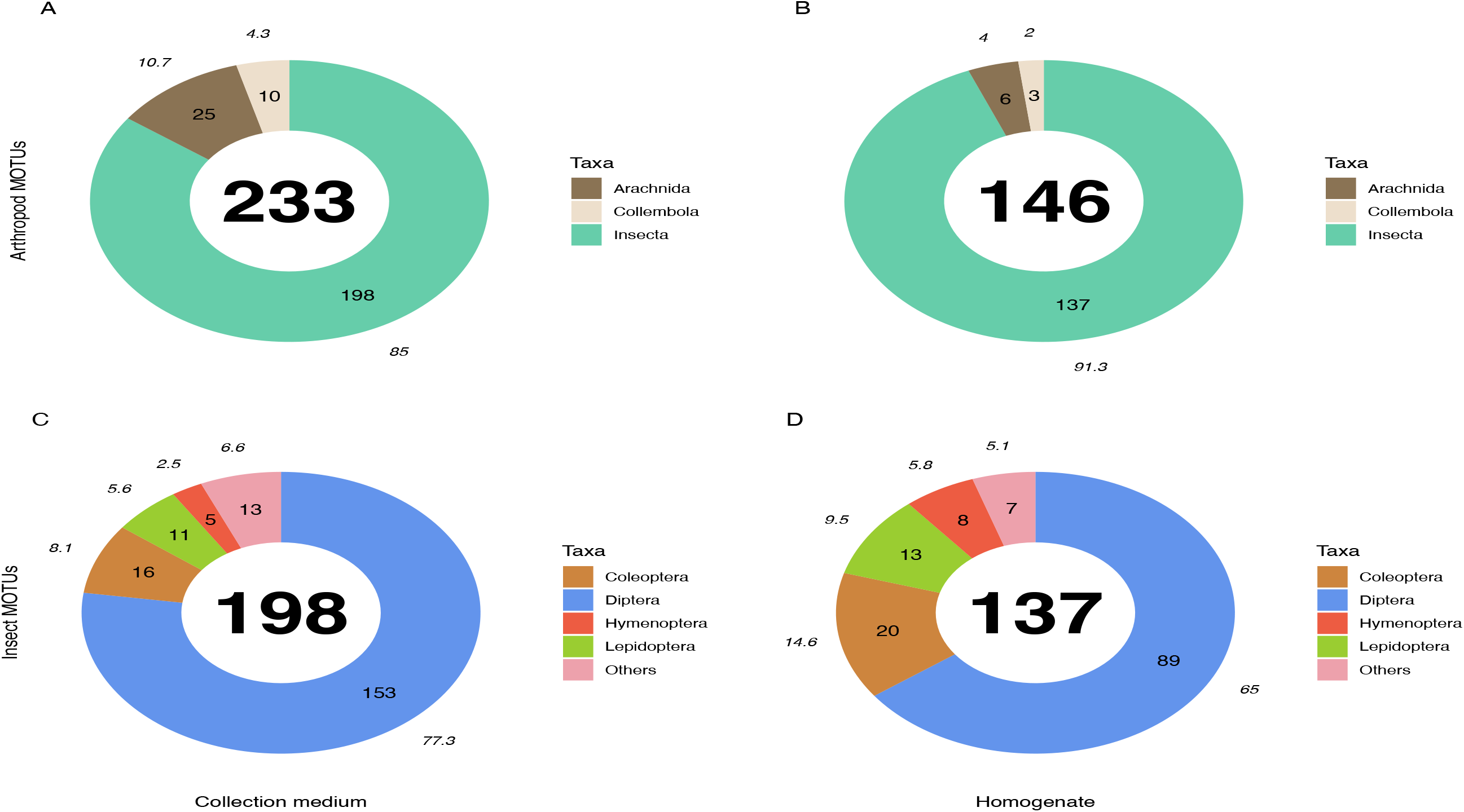
Taxonomic composition (number of MOTUs) of arthropod communities recovered from both homogenate and collection medium metabarcoding of Malaise trap samples. Taxonomic composition (% (italics) and absolute numbers are reported) of MOTUs retrieved from collection medium metabarcoding (**A**, **C**) and homogenate metabarcoding (**B**, **D**) of the same Malaise trap samples. **A**& **B** show the number of MOTUS per Arthropoda classes recovered from homogenate and collection medium respectively. **C**& **D** show the four insect orders with the highest number of MOTUs for homogenate and collection medium respectively. Insects included in the “Others” category belong to Neuroptera, Psocodea and Raphidioptera as well as to Ephemeroptera, Mecoptera, Thysanoptera and Trichoptera in collection medium (**C**) and Hemiptera in homogenate (**D**).

The numbers of detected MOTUs for non-insect taxa (*e.g*. Collembola and Arachnida) was significantly higher in collection medium than in homogenate metabarcoding (Pairwise *T*-test: 1–2: *p* = 6.6^e-03^), similar for Diptera (Wilcoxon rank sum-test: 1–2: *p* = 0.15) and the category “other insect orders” (W-test: 1–2: *p* = 1), but significantly lower for Coleoptera (W-test: 1–2: *p* = 3.9^e-03^), Hymenoptera (W-test: 1–2: *p* = 1.9^e-03^) and Lepidoptera (W-test: 1–2: *p* = 1.4^e-02^) (Supplementary Figure 1).

Comparisons of MOTU consensus sequences between collection medium and homogenate metabarcoding gave 71/233 exact MOTU matches (Figure 6A), of which 18 suggesting that DNA from the same individual can genuinely be recovered by both treatments of the same sample. When considering MOTUs that were identified to species level—118/233 for collection medium and 115/146 for homogenate metabarcoding (Figure 3; 6B)—, 40 species were shared between both treatments (Figure 6B). However, only 9 species were recovered by both treatments of the same sample. (Supplementary Table VII).

**Figure 6:**
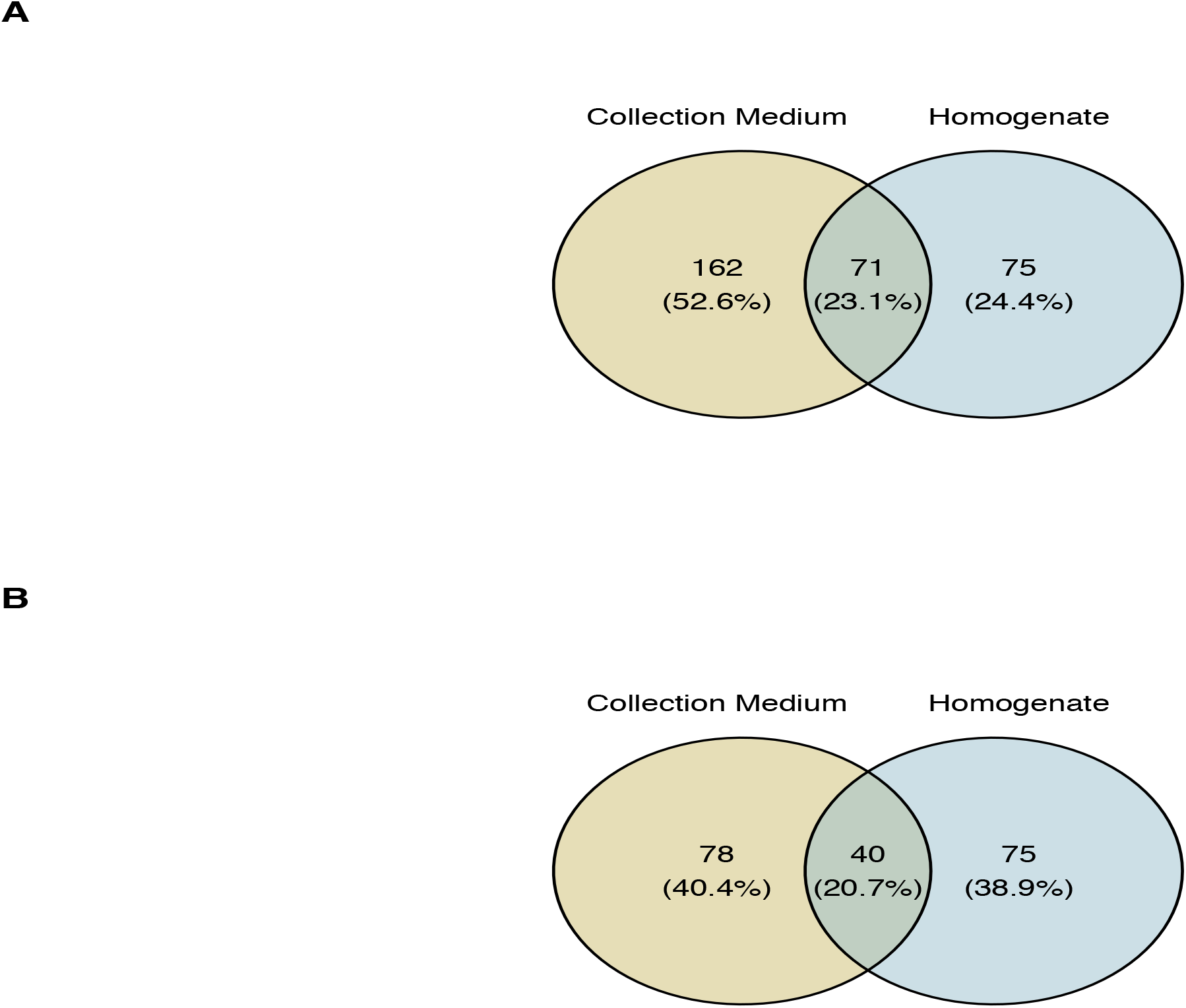
Taxonomic overlap between collection medium and homogenate metabarcoding from Malaise traps. Venn diagram of the total number of MOTUs (**A**) or MOTUs identified to species level (**B**) for homogenate metabarcoding (blue) and collection medium (yellow) of Malaise trap samples. **(A)** 71 MOTUs are shared between collection medium and homogenate metabarcoding, while **(B)** 40 species are shared by both sample types.

We detected no significant change in MOTU richness in collection medium of MT samples among dieback levels (anova: df = 2, *p* = 0.91) or stand types (anova: df = 2, *p* = 0.634) (Figure 7).

**Figure 7:**
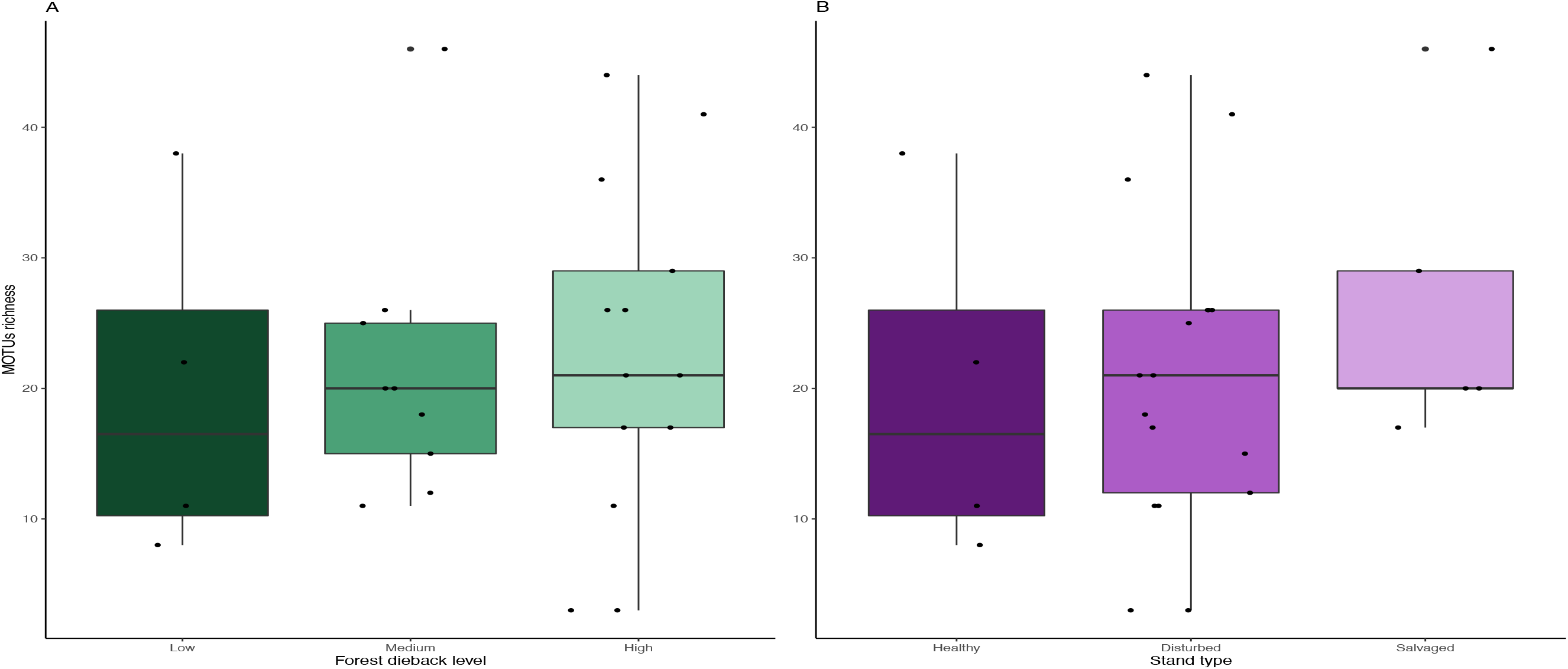
Variation in MOTUs richness across natural and anthropogenic disturbances. Comparison of MOTUs richness recovered from collection medium metabarcoding. Richness variations are tested across (**A**) low, medium and high climate-induced dieback levels and (**B**) between disturbed but unmanaged and salvage-logged plots. Black dots represent samples. No significant differences could be detected with anova tests for both disturbances’ gradients (Dieback level: df = 2, *p* = 0.91; Stand type: df = 2, *p* = 0.634).

## Discussion

### From fieldwork to bioinformatic demultiplexing—technical considerations for collection medium metabarcoding

DNA metabarcoding from bulk samples of arthropods has flourished in the past 10 years, and with it arose many technical considerations from the experimental to the bioinformatic demultiplexing steps (Alberdi *et al*. 2018; Elbrecht *et al*. 2019). Limitations are also being identified for DNA metabarcoding from collection medium and preservative ethanol (Martins *et al*. 2020), but studies remain scarce. Our analyses corroborated the possibility to detect species from collection medium metabarcoding, but the low richness of MOTUs detected in most samples is clearly not representative of the diversity that MTs and WFTs passively collect. Here, we discuss some critical steps that may directly impact EtOH-based metabarcoding results and should be further investigated to test the efficiency and robustness of the approach prior standardization and ecological applications.

Considering field conditions, one factor that could explain the relatively low number of MOTUs detected is the fact that trap jars are often set in clearings or open canopies, hence exposed to warm temperatures and direct UV-light likely accelerating DNA degradation in the field. In addition, drowned organisms also passively release water by osmolarity and dilute the collection medium, which might reduce its preservative capacity when great biomass is accumulated in the trap and also increase risks of DNA hydrolyses (Jo *et al*. 2019). In addition, our samples were collected after one-month in the field which could have led to greater DNA degradation and explain the relatively low MOTU detection rate. Therefore, it is advisable to replace the bottles of malaise traps every one to two weeks maximum to minimize DNA degradation and optimize passive diffusion (Martins *et al*. 2019), with sample storage (or pre-processed filters in case of storage shortage) at −20°C (Yamanaka *et al*. 2016).

The chemical composition of the collection medium may also directly play a critical role on the preservation of extracellular free DNA (*i.e*. DNA molecules passively released by organisms into the collection medium). To avoid DNA hydrolysis (Jo *et al*. 2019), water should be minimized in collection media. However, the substitution of water by ethanol in WFTs leads to higher evaporation rates and costs, increased attractiveness to some insects and subsequent sampling biases (Bouget *et al*. 2009). Furthermore, WFTs are by design exposed to rainfall due to their wide opening on the collector and thus prone to increased water content, the volume of which is limited by small holes drilled on the container to avoid overflowing, but leading to liquid loss and extracellular DNA dilution. Alternative collection media include NaCl solution, either pure or mixed with MPG (Milián-Garcίa *et al*. 2020). Salted water has been shown to be cost-effective for monitoring Coleoptera (Young *et al*. 2020) but may further degrade DNA in traps focusing on soft-bodied taxa with quicker passive DNA diffusion, although this is untested by metabarcoding. Pure MPG collection medium is a good preservative (Stoeckle *et al*. 2010; Höfer *et al*. 2015; Nakamura *et al*. 2020; Martoni *et al*. 2021) but its high viscosity (Martoni *et al*. 2021) might facilitate individual escapes due to increased floatability (McCravy *et al*. 2007), it also might coat free DNA molecules and/or clog the filter membrane (as experienced when filtering 100 mL of collection media containing 50% MPG), all of which may reduce DNA recovery.

During wet-lab processing, several steps may also impact DNA recovery. First, the choice of filters used for DNA isolation may be critical as capture efficiency depends on DNA polarity, which may be affected by the chemical composition of the collection medium. Based on Li *et al*. (2018) results on eDNA filtered from water, we chose mixed-ester cellulose filters for our collection media samples. Other studies successfully captured DNA with nitrate filters from preservative ethanol (Milián-Garcίa *et al*. 2020; Young *et al*. 2020), with an additional grinding step of the membrane to increase lysis efficiency (Kirse *et al*. 2022). However, collection medium might also accumulate inhibitors released from arthropods (Boncristiani *et al*. 2011; Linard *et al*. 2016) or from external by-catches (*i.e*. leaves or pine needles releasing pigments and terpenes (Tang *et al*. 2011), molluscs or worms with high polysaccharide contents), that are likely retained by the filter. Similar inhibition and DNA purity issues have been reported for non-destructive lysis buffer extractions (Kirse *et al*. 2022). Thus, questions on DNA-binding and polarity, filter capture and retention capacities, or pore size and fluidity/clogging remain and should be further explored to evaluate the impact on both free DNA and potential inhibitors yielded from different EtOH-based solutions (and non-destructive alternatives more generally; Kirse *et al*. 2022).

Primer efficiency is a second key factor (Martoni *et al*. 2022) and our analyses showed a lower MOTU richness recovered with Uni-Minibar primers compared to the commonly used 313-bp COI fragment amplified by the mlCOIintF/jgHCO2198 primer set (Leray *et al*. 2013; Geller *et al*. 2013). Unfortunately, PCR targeting 313-bp amplicons failed on collection media. Moreover, this COI fragment does not overlap with the 127-bp Uni-Minibar fragment amplified, making MOTU comparisons through alignments impossible (Elbrecht *et al*. 2019). Thus, to compare the efficiency of DNA metabarcoding between treatments (MT homogenate metabarcoding *vs*. MT collection medium metabarcoding) we had to use the Uni-Minibar primers’ amplicon for homogenate. As diversity recovered was significantly lower with the Uni-Minibar primers than with longer amplicons allowing increased resolution (Figure 4, Supplementary Figure 1), it is likely that similar amplification and identification biases has been obtained from metabarcoding the collection media.

Lastly, bioinformatic processing is also instrumental to determine MOTU diversity. In particular, demultiplexing parameters on filtering MOTUs across different PCR replicates can greatly impact numbers of sequence reads and MOTU retained (Alberdi *et al*. 2018). Regardless of the type of trap (WFT and MT), the use of a more conservative retention (MOTUs present in at least two PCRs) allowed a drastic reduction of unknown sequences and chimeras, untargeted organisms, or contaminants, but did not lead to an important decrease in identified and plausible species. It also suggests that sequencing depths allocated to sequence species present in the samples was diminished, further influencing the poor results on our MOTU recovery.

### Community analyses and terrestrial insect monitoring from collection medium metabarcoding of Malaise trap samples

Accurate species identification is crucial to ecological analyses, to unravel species biology and the functions they may have in their respective environments (Tautz *et al*. 2003). In environmental genomics, community analyses based on metabarcoding rely on DNA reference libraries to identify species. While metabarcoding collection medium allows for the preservation of voucher specimens for morphological validation, it remains important to assess whether this molecular approach can reliably inform insect communities.

Here, taxonomic assignment at species level was the lowest for Diptera (51%), Arachnida (16%) and Collembola (10%). This may be explained by the fact that these groups are highly diverse and notoriously difficult to identify based morphological criteria, or are poorly covered in DNA barcode reference libraries (Morinière *et al*. 2019; Sire *et al*. 2022). However, thanks to the recent DNA barcoding efforts to cover the fauna of Germany it is possible to identify a relatively large proportion of the Central and Western European dipteran fauna (Morinière *et al*. 2019). It is also of note that the short length of the amplicon targeted here (127 bp) reduces taxonomic resolution (Hajibabaei *et al*. 2006; Meusnier *et al*. 2008; Elbrecht *et al*. 2019). Interestingly, we found that the insect communities characterised with collection medium metabarcoding and homogenate metabarcoding for the same MT samples were overall dissimilar, with only 71 MOTUs or 40 identified species shared between collection medium and homogenate metabarcoding (Figure 6). Comparisons at class and order levels also suggest that collection medium metabarcoding slightly differs from homogenate metabarcoding.

These discrepancies of results between collection medium and homogenate metabarcoding of a MT sample are in line with previous reports showing dissimilar communities, especially the higher detection of soft-bodied (poorly sclerotized) arthropods like Arachnida and Collembola and a large dipteran diversity, or an under-detection of Coleoptera in collection medium (Marquina *et al*. 2019; Kirse *et al*. 2022; Chimeno *et al*. 2022b, Martoni *et al*. 2022). As dipterans are a highly diverse and functionally important group of insects (*e.g*. pollinators, decomposers, *etc*.) in forest ecosystems (Mlynarek *et al*. 2018; Chimeno *et al*. 2022a), the use of EtOH–MPG collection medium metabarcoding could improve our understanding of their ecological role at the community level for environmental assessment. In contrast, we show an unusually low detection of Hymenoptera MOTUs, which is likely caused by the low affinity of Uni-Minibar primers toward this order (Yu *et al*. 2012; Brandon-Mong *et al*. 2015). Collection medium metabarcoding is therefore unlikely to strictly substitute homogenate metabarcoding (Marquina *et al*. 2019). Running both treatments in parallel could instead enrich biodiversity surveys and broaden our understanding of trophic assemblages. In particular, medium-based metabarcoding may outperform bulk-based approaches for the detection of prey DNA that is regurgitated or defecated by captured organisms at the time of death, or for the recovery of DNA from pollen and fungi spores brought by the arthropods falling in the traps. The caveats of homogenate metabarcoding remains the loss of voucher specimens that impedes subsequent morphological studies, DNA barcoding of individuals and collection storing (Marquina *et al*. 2019). This may also hinder the transition for metabarcoding-based biodiversity survey if sample preservation is legally mandatory in official biomonitoring programs (Martins *et al*. 2019). Interestingly, this problem may not apply to other types of samples as in surveys of freshwater organisms, similar taxonomic recoveries were found by metabarcoding EtOH preservative and homogenates (Hajibabaei *et al*. 2012; Zizka *et al*. 2018). As there are no standardized laboratory procedures, comparisons between sample types and studies remain difficult. However, these discrepancies in species recovery patterns may reflect the differences among sample types and highlight the need to assess sample provenance and clarity for reliable comparisons (Box 1; (Martins *et al*. 2019, 2020).

Although metabarcoding collection medium or homogenate documented different arthropod communities, both methods may have comparable value for monitoring the response of species assemblages to environmental changes—in our case the response of arthropods to forest dieback gradient induced by droughts and associated forest management. No response could be detected in terms of MOTUs richness across the three levels of climate-induced forest dieback intensity, nor between the three various stand types. This result is similar to a previous broader study that included the samples analysed here (Sire *et al*. 2022). However, the relatively low success of MOTU recovery impedes further analyses on community changes to evaluate ecological and functional responses as investigated from homogenate metabarcoding of these samples using Leray/Geller primers (Sire *et al*., 2022). Interestingly, Chimeno *et al*. (2022b) showed that Malaise trap communities across their two treatments (*i.e*. preservative EtOH *vs*. homogenate metabarcoding) were dissimilar and highlighted that communities recovered from EtOH-based metabarcoding differed in their composition and response to environmental changes from those recovered from homogenate metabarcoding. This is in contrast with previous studies highlighting the potential to monitor freshwater ecosystems (Zizka *et al*. 2018; Martins *et al*. 2019, 2020; Persaud *et al*. 2021) or population genetics (Couton *et al*. 2021) with EtOH-based metabarcoding as a potential replacement for homogenate metabarcoding.

## Conclusion

Our study brings another example of the use of non-destructive collection/preservation medium-based metabarcoding for the survey of terrestrial arthropods. Our use of collection medium metabarcoding informed communities that differ from those obtained using homogenate metabarcoding and complemented that approach, possibly through increased detection of small and soft-bodied organisms or ingested DNA released by predators. Analyzing the metagenome of collection/preservation medium takes metabarcoding away from ideal experimental conditions and we expect it to be much impacted by fieldwork conditions (DNA degradation, inhibitors, collection medium composition), laboratory processes (storage and contaminants, DNA filtering and extraction, primer affinity) and data analysis (sequence length, sequencing depth). In that sense, medium-based metabarcoding requires further methodological developments and testing to unlock its full potential—a goal worth pursuing, especially when sampling the poorly known arthropod fauna (Lopez-Vaamonde *et al*. 2019) of biodiversity hotspots where preserving the integrity of specimens is most important for further description and study.

## Supporting information

Supplementary Tables I to VII

## Acknowledgments

We are thankful to Wilfried Heintz, Laurent Burnel, Jérôme Molina and Jérôme Willm for the field work, Sylvie Ladet for her SIG assistance, Carl Moliard and Guilhem Parmain for the sorting and morphological identification of the Coleoptera from WFTs. We also want to thank Florent Figon and Sophie Van Meyel for helpful discussions on the manuscript. Part of this work has been carried out with the technical support of the Genomic Facilities (PST ‘Analyse des Systèmes Biologiques’) at university of Tours.

## Author Contributions

- Lucas Sire conceived and designed the experiment, performed the experiment, analysed the data, prepared figures and/or tables, authored and reviewed drafts of the paper.
- Paul Schmidt Yáñez analysed the data, reviewed drafts of the paper.
- Annie Bézier performed the experiment, reviewed drafts of the paper.
- Béatrice Courtial performed the experiment, reviewed drafts of the paper.
- Susan Mbedi performed the experiment, reviewed drafts of the paper.
- Sarah Sparmann performed the experiment, reviewed drafts of the paper.
- Laurent Larrieu conceived and designed the experiment, reviewed drafts of the paper.
- Rodolphe Rougerie reviewed drafts of the paper.
- Christophe Bouget conceived and designed the experiment, reviewed drafts of the paper.
- Michael T. Monaghan reviewed drafts of the paper.
- Elisabeth A. Herniou conceived and designed the experiment, reviewed drafts of the paper.
- Carlos Lopez-Vaamonde conceived and designed the experiment, reviewed drafts of the paper.

## Data Availability

The raw sequencing data are available at NCBI in the following BioProject: PRJNA927244 : CLIMTREE_EtOH.

## Boxes, Figures, Tables and Supplementary

**<u>Box 1:</u> Terminology and sample types in non-destructive metabarcoding: differences between collection medium and preservative ethanol.**

The exploratory nature of non-destructive metabarcoding from various liquids makes comparison difficult, especially due to the type of samples used and the aquatic or terrestrial origin of the targeted arthropod communities (Zizka *et al*. 2018; Erdozain *et al*. 2019; Marquina *et al*. 2019; Martins *et al*. 2019, 2020; Zenker *et al*. 2019; Milián-Garcia *et al*. 2020; Young *et al*. 2020; Zenker *et al*. 2020; Persaud *et al*. 2021; Wang *et al*. 2021b, Chimeno *et al*. 2022b). In most of these studies, the word used to describe the sample type is “preservative ethanol”. However, sample type and liquid “clarity”, or “dirtiness” as called by Martins *et al*. (2019), can be quite different according to facultative pre-processing steps, or the arthropod community targeted, and this may significantly alter the information recovered from metabarcoding. Therefore, we propose a terminology that precisely reflects the sample type used (Figure B-1).

To illustrate our point, terrestrial arthropods and especially insects are often sampled with passive-sampling trapping methods like Malaise traps (MT) or window-flight traps (WFT). Both collect insects directly within a trapping liquid which stays in the field during a variable time period (*e.g*. one week to one month). This trapping liquid from which insects are filtered out without further processing is what we call “**collection medium**”, and is the liquid type used by some studies like Marquina *et al*. (2019), Milián-Garcia *et al*. (2020), Young *et al*. (2020) or Kirse *et al*. (2022). Filtered insects can then be morphologically sorted (Young *et al*. 2020), individually barcoded or processed via metabarcoding from DNA extraction from insects that have been grinded-down to powder (Yu *et al*. 2012; Sire *et al*. 2022) that we define here similarly to Marquina *et al*. (2019) as **homogenate metabarcoding**. Alternatively, filtered insects can also be placed in fresh ethanol during a variable time period for voucher preservation and storage, and can be filtered out again from this ethanol for further morphological or molecular analyses. The liquid recovered after this second filtration of insects out of ethanol gives a second sample type that we call here “**preservative ethanol**” and that we consider different from collection medium (Figure B-1). Currently, this sample type matches the sample description of most of the studies on ethanol-based metabarcoding (Shokralla *et al*. 2010; Hajibabaei *et al*. 2012; Linard *et al*. 2016; Zizka *et al*. 2018; Erdozain *et al*. 2019; Martins *et al*. 2019, 2020; Zenker *et al*. 2020; Persaud *et al*. 2021; Wang *et al*. 2021b; Chimeno *et al*. 2022b).

There are notable differences between the two sample types. First whereas preservative ethanol is—as indicated by its name—pure ethanol (which may vary in titrations), collection medium encompasses various chemical compositions based on pure liquids or mixtures (*e.g*. water, salted water, (monopropylene) glycol, ethanol, ethyl acetate, soap…). Second, collection medium is the dirtiest, as it contains environmental debris and/or arthropod outer-exoskeleton (free-)DNA materials (*e.g*., pollen, dirt, leave debris, fungi spores, ectoparasites…). Collection medium also contains ingested DNA (iDNA) from intestinal and/or gut contents potentially released by regurgitation and/or defecation death reflexes during insect drowning (Marquina *et al*. 2019). In comparison, preservative ethanol is relatively clear and free-DNA mostly derives from passive diffusion of the dead arthropods present in the bottle. Of note, the clear/dirty qualification is not binary but rather a continuous gradient that depends of the targeted communities, whether organisms are alive as they get into the liquid used for DNA extraction, or according to the sample’s surrounding environment and its time spent in the field (Figure B-1). It follows that samples of freshwater communities from the previously listed studies are more similar to preservative ethanol than to collection medium, for three reasons: (*i*) arthropods are less likely to carry outer-exoskeleton DNA material as evolving in aquatic environments, after kick-net sampling—that can be extremely dirty—arthropods are often sorted-out of environmental debris prior to ethanol transfer, (*iii*) life-status prior ethanol transfer is often uncertain (except for live transfer described in Linard *et al*. (2016)), reducing their potentiality to yield iDNA from similar death reflexes as for terrestrial insects. We acknowledge that these points can be nuanced for kick-net samples (*e.g*. caddisfly larva cases result in both organic and/or non-organic inputs, kick-net sorting is not compulsory (Pereira-da-Conceicoa *et al*. 2020), *etc*) and each case should be explicitly described for further comparisons and robustness.

Information on insect sampling is therefore crucial to correctly categorize the processed samples. Thus, we recommend to distinguish collection medium from preservative ethanol as described above to facilitate cross comparisons between studies and recommend to mention whether arthropods are alive and pre-sorted prior to be transferred in preservative ethanol.

**Figure B-1:**
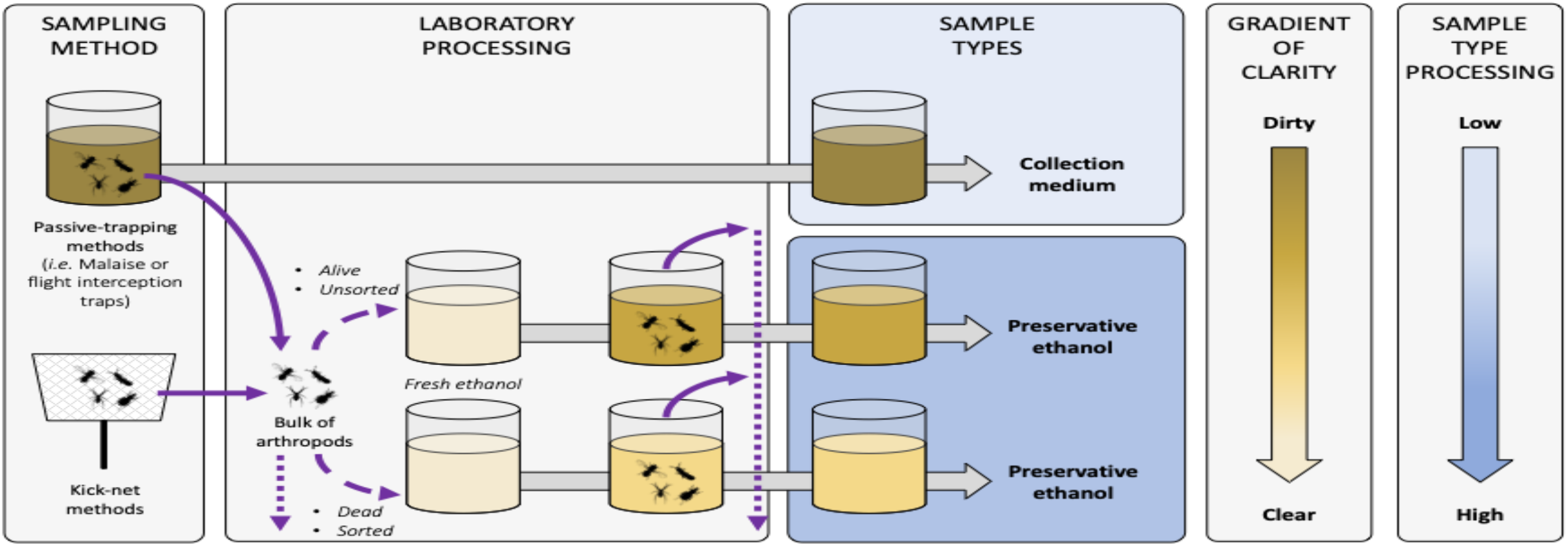
Terminology and description of sample types for metabarcoding from trapping liquids. Diagram representing the sample types that can be used when metabarcoding collection or preservative media. Solid and dashed violet arrows represent arthropods transferred in and out of liquids, respectively. Arthropod live-status (*i.e*. dead or alive) and sample condition (*i.e*. sorted / unsorted) are listed as factors influencing the clarity of the sample. Dotted violet arrows represent arthropod post-processing potentialities (*i.e*. morphological sorting, DNA barcoding or metabarcoding, storing…). Grey arrows represent time processing that can be variable before sample sequencing. Sample shades of yellow represent the clarity of the liquid sample, with the darker the dirtier according to the gradient of clarity on the right, and with fresh ethanol in light yellow as the clearest and equivalent to a blank control. Sample types boxes are coloured according to the level of sample processing and manipulation post-sampling according to the shaded blue gradient on the right, with light blue the lowest and dark blue the highest amount of sample handling, respectively.

**Table I:** Summary of the MOTUs recovery success for each trapping method and sample type analysis. WFT: window-flight trap.

| Sample type | # samples processed | # samples recovered (%) | Min (M)OTUs per recovered sample | Max (M)OTUs per recovered sample | Mean (M)OTUs per recovered sample | Total (M)OTUs recovered |
| --- | --- | --- | --- | --- | --- | --- |
| Malaise trap (*collection media*) | 27 | 26 (96%) | 3 | 46 | 21.81 | 233 |
| Malaise trap (*homogenate*) | 27 | 10 (37%) | 17 | 50 | 32.4 | 146 |
| WFT (*collection media*) | 53 | 37 (70%) | 1 | 47 | 2.06 | 77 |
| WFT (*morphology*) | 53 | 53 (100%) | 22 | 82 | 54.43 | 389 |

**Supplementary Figure 1:**
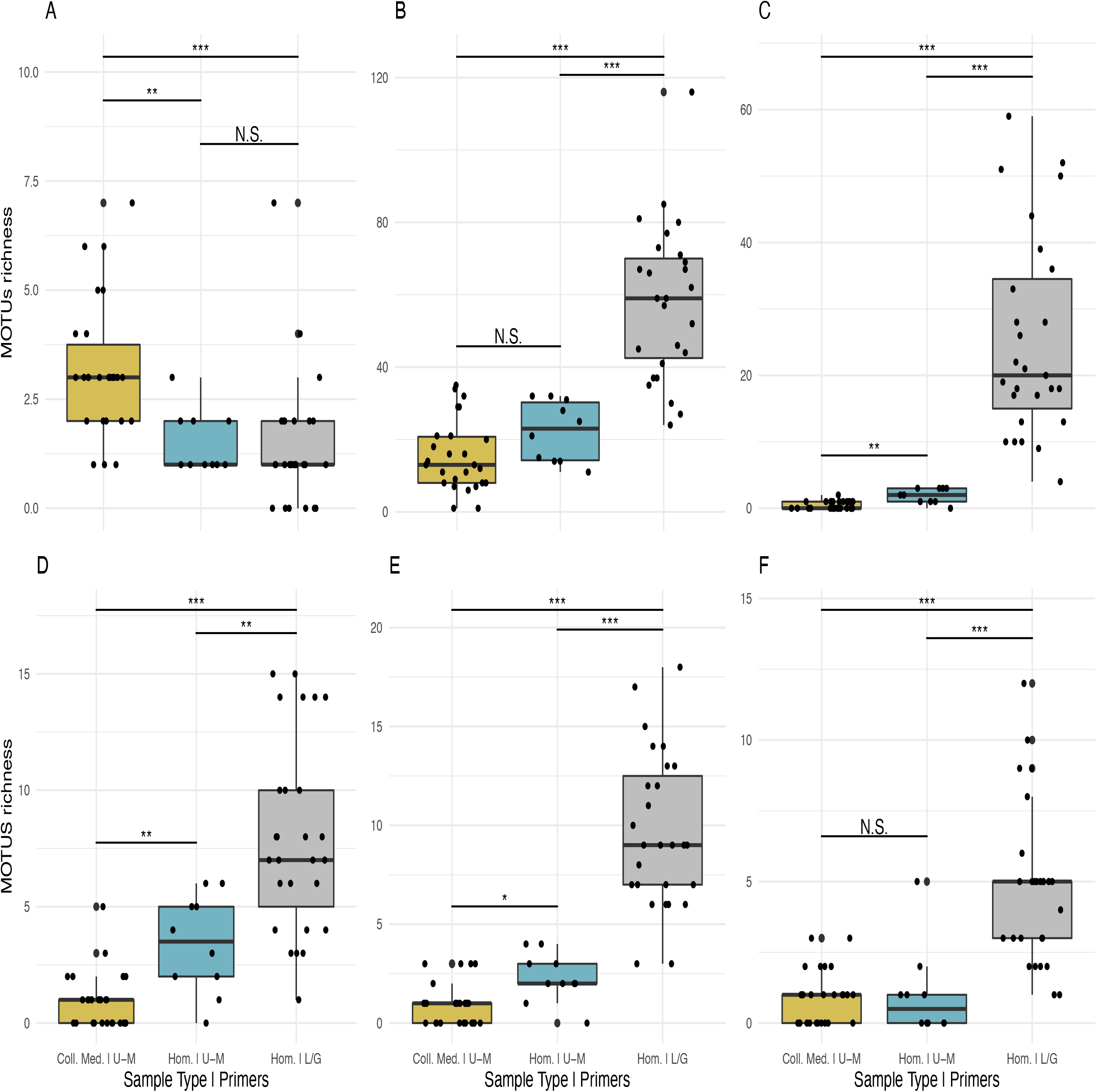
Comparison of MOTU richness for different insect taxa recovered from Malaise traps using metabarcoding of collection medium (Coll. Med.) or homogenate (Hom.) with the Uni-Minibar (U-M) or Leray/Geller (L/G) primer sets. Boxplot of MOTU count for collection medium (yellow; 1) or homogenate metabarcoding (blue; 2) with Uni-Minibar primer set or from homogenate metabarcoding using mlCOIintF/jgHCO2198 primer set (gray; 3) of the same Malaise trap samples. Black dots represent samples considered after demultiplexing and data curation. Significant differences adjusted with Bonferroni correction are highlighted with ‘*’ and ‘N.S.’ stands as non-significant. Studied taxa are: (**A**) non Insecta (*i.e*. Arachnida and Collembola) (Pairwise *T*-test: 1–2: *p* = 6.6^e-03^; 1–3: *p* = 7.4^e-05^; 2–3: *p* = 1); (**B**) Diptera (Wilcoxon rank sum-test: 1–2: *p* = 0.15; 1–3: *p* = 6.0^e-09^; 2–3: *p* = 8.3^e-05^); (**C**) Hymenoptera (W-test: 1–2: *p* = 1.9^e-03^; 1–3: *p* = 7.6^e-10^; 2–3: *p* = 1.2^e-05^); (**D**) Coleoptera (W-test: 1–2: *p* = 3.9^e-03^; 1–3: *p* = 6.6^e-09^; 2–3: *p* = 4.2^e-03^); (E) Lepidoptera (W-test: 1–2: *p* = 1.4^e-02^; 1–3: *p* = 1.5^e-09^; 2–3: *p* = 3.1^e-05^); (**F**) other Insecta orders grouped (W-test: 1–2: *p* = 1; 1–3: *p* = 7.7^e-08^; 2–3: *p* = 3.5^e-04^).

